# SKP2 knockout in Rb1/p53 deficient mouse models of osteosarcoma induces immune infiltration and drives a transcriptional program with a favorable prognosis

**DOI:** 10.1101/2023.05.09.540053

**Authors:** Alexander Ferrena, Jichuan Wang, Ranxin Zhang, Burcu Karadal-Ferrena, Waleed Al-Hardan, Swapnil Singh, Hasibagan Borjihan, Edward Schwartz, Hongling Zhao, Rui Yang, David Geller, Bang Hoang, Deyou Zheng

**Affiliations:** Institute for Clinical and Translational Research, Albert Einstein College of Medicine, Bronx, NY, USA; Department of Genetics, Albert Einstein College of Medicine, Bronx, NY, USA; Department of Orthopedic Surgery, Montefiore Medical Center, Albert Einstein College of Medicine, Bronx, NY, USA; Department of Pathology, Albert Einstein College of Medicine, Bronx, NY, USA; Department of Oncology, Albert Einstein College of Medicine, Bronx, NY, USA; Department of Medicine, Albert Einstein College of Medicine, Bronx, NY, USA; Department of Molecular Pharmacology, Albert Einstein College of Medicine, Bronx, NY, USA; Department of Developmental & Molecular Biology, Albert Einstein College of Medicine, Bronx, NY, USA; Department of Neurology, Albert Einstein College of Medicine, Bronx, NY, USA; Department of Neuroscience, Albert Einstein College of Medicine, Bronx, NY, USA

**Keywords:** Osteosarcoma, SKP2, RNA-seq, TME, Immune infiltration

## Abstract

**Purpose:** Osteosarcoma (OS) is an aggressive bone malignancy with a poor prognosis. One putative proto-oncogene in OS is *SKP2*, encoding a substrate recognition factor of the SCF E3 ubiquitin ligase. We previously demonstrated that *SKP2* knockout in murine OS improved survival and delayed tumorigenesis. Here we aim to define the SKP2 drives transcriptional program and its clinical implication in OS.

**Experimental Design:** We performed RNA-sequencing (RNA-seq) on tumors from a transgenic OS mouse model with conditional *Trp53* and *Rb1* knockouts in the osteoblast lineage (“DKO”: Osx1-Cre;Rb1^lox/lox^;p53^lox/lox^) and a triple-knockout model with additional *Skp2* germline knockout (“TKO”: Osx1-Cre;Rb1^lox/lox^;p53^lox/lox^;SKP2^−/−^). We validated our RNA-seq findings using qPCR and immunohistochemistry. To investigate the clinical implications of our results, we analyzed a human OS patient cohort (“NCI-TARGET OS”) with RNA-seq and clinical data.

**Results:** We found large differences in gene expression after SKP2 knockout. Strikingly, we observed increased expression of genes related to immune microenvironment infiltration in TKO tumors. We observed significant increases in signature genes for macrophages and to a lesser extent, T cells, B cells and vascular cells. We also uncovered a set of relevant transcription factors that may mediate the changes. In OS patient cohorts, high expression of genes upregulated in TKO was correlated with favorable overall survival, which was largely explained by the macrophage gene signatures. This relationship was further supported by our finding that SKP2 expression was negatively correlated with macrophage infiltration in the NCI-TARGET OS and the TCGA Sarcoma cohort.

**Conclusion:** Our findings indicate that SKP2 may mediate immune exclusion from the OS tumor microenvironment, suggesting that SKP2 modulation in OS may induce anti-tumor immune activation.

**Translational Relevance:** Osteosarcoma (OS) is an aggressive bone malignancy. Standard care treatment involving chemotherapy and surgery remains unchanged for decades. OS prognosis remains poor and targeted therapies are critically needed. Immunotherapy failed in clinical trials despite extensive genomic instability, suggesting OS tumors employ active immune exclusion. One putative oncogene in OS is *SKP2*, which activates cell proliferation. *SKP2* is correlated with worse prognosis in patients and knockout of *SKP2* improved survival in murine OS. Here, we performed comparative transcriptomic analysis between transgenic OS murine models with *SKP2* knockout and controls. We showed that *SKP2* knockout dramatic increased immune gene expression largely due to macrophage, and to a lesser extent lymphocytes, infiltration. Interestingly, we found that the increased gene program uncovered from our *SKP2* knockout model was correlated with improved survival in OS patients. Our findings indicate a completely new function for *SKP2* in mediating immune exclusion in sarcoma and suggest that *SKP2* inhibition may lead to improved immune activation and potential treatment.

## Introduction

Osteosarcoma (OS) is a cancer of bone tissue defined by mesenchymal cell histology and deposition of bone matrix^1, 2^. OS is the most common pediatric bone malignancy with annual US incidence of 3.1 cases per million^3^. Disease progression commonly involves metastasis to the lungs^4^. For cases that are non-metastatic at diagnosis, 5-year survival is 70% while for metastatic cases at presentation, comprising 15 to 25% of incidents, the rate is 30%^4, 5^. The cell origin in OS is unknown but is thought to be cells in the osteoblast lineage ranging from mesenchymal stem cells (MSCs) to pre-osteoblasts polarized to the osteoblast lineage^6^. Osteoblast differentiation from multipotent MSC progenitors requires the expression of lineage-specific transcription factors, including *Osterix* (Osx)^7^. Genetically, OS is complex and involves many copy number alterations, but the two most frequent mutations are loss of the tumor suppressors *TP53* and *RB1* ^8–12^. *In vivo* investigation in preclinical murine models revealed that ablation of *Trp53* and *Rb1* by *Osx*-driven Cre-Lox genetic recombination generates bone tumors that strongly resemble human OS in histology, cytogenetics, and metastatic progression^13^. Our recent analysis of clinical genomic datasets revealed that *TP53* and *RB1* were respectively the first and second most frequently mutated genes in OS and furthermore were often co-mutated^14^.

S-phase Kinase Associated Protein 2 (*SKP2*) codes for an F-box substrate recognition factor of the Skp1-Cullin1-F-box (SCF) E3 ubiquitin ligase complex. Its known functions include activation of the cell cycle, cell growth, and migration, along with repression of apoptosis, via canonical proteolytic K48 ubiquitination and non-proteolytic K63 ubiquitination^15^. Two key targets for proteolytic ubiquitination by SCF^SKP2^ are the cyclin-dependent kinase inhibitors p21 and p27, allowing cell cycle progression^16, 17^. In mouse models of pituitary and prostate cancers in the context of *Rb1* and *Trp53* loss, *Skp2* knockout was shown to be synthetic-lethal, permanently blocking tumorigenesis in a p27-mediated manner^18^. Mechanistically, *SKP2* inactivation in the context of *Rb1* deficiency did not block proliferation, but instead resulted in p27-mediated apoptosis^19^. Specifically, *Rb1* deletion induced activation of the transcription factor *E2f1* resulting in proliferative gene expression, but co-deletion of *Rb1* and *Skp2* allowed accumulated p27 to form more Cyclin A-p27 complex, which releases the restraining effects of Cyclin A on *E2f1*, causing *E2f1* “superactivation” and resulting in p53-independent apoptosis and blockage of tumorigenesis^20^. *SKP2* thus represents an attractive targetable oncogene in OS and other cancers driven by *RB1* and *P53* mutations. However, in other contexts such as *Myc*-driven lymphomagenesis, SKP2 inhibition showed little effect^21^.

Pharmacologic inhibitors of SKP2 include Pevonedistat, an inhibitor of neddylation, a key upstream post-translation modification required for SCF^SKP2^ and other cullin-RING E3 ligases^22^. Other, more specific inhibitors include “C1”, which blocks the SKP2-p27 interaction, and “C25”, which blocks SKP2’s interaction with SKP1 that is required for SCF’s E3 ubiquitylation activity^23, 24^. Our previous work has demonstrated that pharmacologic inhibition of SKP2 reduced invasion and lung metastasis in patient-derived xenograft models of OS^25^. Additionally, blocking the proteolytic ubiquitination of p27 by SKP2 *in vivo* in a transgenic mouse model of OS with both genetic and pharmacologic methods promoted survival and induced apoptosis and cell cycle arrest^14^. Most recently, we have shown that genetic knockout and pharmacologic inhibition of SKP2 *in vivo* further improved survival and promoted tumor apoptosis in the mouse DKO OS model^26^. However, unlike in models of pituitary and prostate cancers, OS tumors with SKP2 knockout continue to develop and progress, suggesting undescribed context-specific resistance mechanisms. Additionally, despite the importance of SKP2 to cell proliferation and cancer, to our knowledge no data on the impact of SKP2 knockout on the transcriptome has been reported.

Immune and stromal infiltration into the tumor microenvironment (TME) is widely recognized as a key characteristic of solid tumor malignancies. In OS, single-cell RNA-seq studies have reported a diverse ecosystem of infiltrating non-malignant cell populations within OS tumors^27, 28^. While OS frequently presents with genomic instability resulting in neoantigens, OS tumors were found to employ mechanisms of lymphocyte exclusion, and checkpoint inhibitors failed to show clinical benefits^29–31^. Macrophages are consistently observed in the TME of many solid malignancies including OS^27, 28^. Many reports have shown that macrophages contribute to cancer disease progression and poor prognosis, for example via promoting pro-metastatic cancer cell intravasation in breast cancer in a manner that may be potentiated by standard-of-care chemotherapies^32, 33^. Conversely, macrophages can also play roles in anti-tumor immunity, including recruitment of cytotoxic T lymphocytes during pro-inflammatory “M1” polarization, and direct cytotoxicity including by macrophage phagocytosis of malignant cells^34, 35^. In OS, macrophage infiltration has been shown to be correlated with improved metastasis-free survival^36^. Several recent pre-clinical reports have described success with macrophage-based immunotherapy. In one such report, combined pharmacologic blockade of CD47 and GD2, two immunomodulatory “don’t eat me” signals which block phagocytosis, was able to reduce tumor burden and extend survival in orthotopic patient-derived xenograft models of OS, along with other cancer types including neuroblastoma and small cell lung cancer^37^. Another study demonstrated that the L-amino acid transporter 2 (*LAT2*) is capable of upregulating mTOR and downstream upregulation of c-Myc and Cd47, and that *LAT2* inhibition was able to enhance macrophage infiltration and phagocytosis of tumor cells and sensitize OS tumor cells to doxorubicin. However, not all prior reports consistently show macrophages portend a positive prognosis in OS^38^. Taken together, the literature indicates that macrophages are a complex, context-dependent component of the TME that may be activated for anti-tumor activity in OS and other malignancies.

Here, we set out to study the transcriptional regulatory roles of SKP2 in OS using murine models. As transgenic mouse models are immunologically intact and give rise to autochthonous tumors, they more closely resemble clinical presentation of OS than cell culture or transplanted tumor models. We compared OS models with germline SKP2 knockout and control using RNA-seq and bioinformatics analysis. Surprisingly, we found that SKP2 knockout drove immune cell (T cell and macrophage) recruitment to the OS TME. Additionally, genes with increased expression upon SKP2 knockout correlated with a significant survival benefit in OS patients in a manner likely driven by macrophage infiltration. Moreover, SKP2 was negatively correlated with macrophage infiltration in OS and soft tissue sarcoma patients. Thus, for the first time, we have uncovered evidence that SKP2 may act as a mediator of immune exclusion from the OS tumor microenvironment.

## Methods

### Establishment of animal models

Osx1-Cre mice, Rb1^lox/lox^ mice, Trp53^lox/lox^ mice, and Skp2^−/−^ mice were described previously^13, 14, 39^. All mice used for experiments are on FVB, C57BL6J, and 129Sv hybrid backgrounds. First, Rb1^lox/lox^ mice were crossed with Trp53^lox/lox^ mice to generate Trp53^lox/lox^, Rb1^lox/lox^, mice, and were further crossed with Osx1-Cre mice to generate Osx1-Cre; Trp53^lox/lox^, Rb1^lox/lox^ mice. The Skp2^−/−^ mice were crossed with Osx1-Cre;Trp53^lox/lox^,Rb1^lox/lox^ mice to generate Osx1-Cre;Rb1^lox/lox^;Trp53^lox/lox^; Skp2^−/−^ mice. Animals were maintained under a pathogen-free condition in the Albert Einstein College of Medicine animal facility. Animal experimental protocols were reviewed and approved by Einstein Animal Care and Use Committee (#20180401), conforming to accepted standards of humane animal care. The tumor diameter was measured using a caliper every three days, and the relative tumor volume was calculated by the following formula: (length x width^2^) x 0.526. Tumors were resected when the tumor reached volume reached approximately 500 mm^3^.

### RNA-sequencing

Mouse tumor tissues from DKO or TKO animals were harvested and processed for RNA-seq. Approximately 100 mg of fresh tumor tissue was collected and processed using the RNeasy Mini Kit (#74104 Qiagen) for RNA extraction following the manufacturer’s protocol. Samples were first quantified and then evaluated for RNA integrity, degradation, and contamination. Next, the rRNA-depleted RNA sample was enriched using oligo(dT) beads and randomly fragmented. This was then followed by cDNA synthesis using random hexamers and reverse transcriptase. After first-strand synthesis, a custom second-strand synthesis buffer (Illumina) was added with dNTPs, RNase H, and *E. coli* polymerase I to generate the second strand by nick translation. The final cDNA library was completed following a round of purification, terminal repair, A-tailing, ligation of sequencing adapters, size selection, and PCR enrichment. The library concentration was quantified using a Qubit 2.0 fluorometer (Life Technologies). The insert size was checked by an Agilent 2100 Bioanalyzer and then further quantified by qPCR. Libraries were then sequenced using an Illumina HiSeq platform. Raw data were processed by the Illumina pipeline, and the resulting RNA-seq reads were mapped to the mouse reference genome (mm10) using the STAR software (v2.6.1) with default parameters ^40^.

### Differential expression analysis

The R package DESeq2 (v1.35.2) was used to perform differential expression analysis^41^. Thresholds for significant differential expression were set to Log2FoldChange > 1 (or < −1) and adjusted P value < 0.05. For overrepresentation analysis (ORA) of significantly differentially expressed genes (DEGs), the generalized ORA function enricher() using Fisher exact test and default parameters from the ClusterProfiler (v4.4.4) package was used to test enrichment of the DEGs in pathways which we downloaded from the Molecular Signatures Database via the R package msigdbr (v7.5.1)^42, 43^. Customized scripts were used to derive gene set clusters and percent of differentially expressed genes using the enrichment maps generated by the emapplot() function from ClusterProfiler.

### Deconvolution analysis of mouse bulk RNA-seq data

For deconvolution of the TKO and DKO bulk RNA-seq tumor data, the R package mouse Microenvironment Cell Population Counter (mMCP-Counter) was used^44^. Infiltration was compared in TKO vs DKO via two-sample T-test. To analyze cytokine expression, we used a cytokine panel derived from the signature “KEGG CYTOKINE CYTOKINE RECEPTOR INTERACTION” downloaded via msigdbr and removed any receptor genes in the panel. Then, we selected for only differentially expressed cytokines for presentation.

### Macrophage polarization analysis

To study macrophage polarization in the TKO bulk RNA-seq data, we downloaded single-cell RNA-seq data of macrophage polarization states^45^. We performed integration, batch correction, and marker analysis of M0, M1, and M2 states using the R package RISC^46^. Using these markers, we performed a Fisher exact test in R to test if markers of these states were enriched in the TKO upregulated genes.

### Immunohistochemical staining and quantitative PCR analysis

Mouse limb tumor tissues were harvested and fixed with 10% formalin, then underwent decalcification with OSTEOSOFT mild decalcifier-solution for histology (Millipore, #101728) for 24 hours or longer until penetrable with a syringe needle. The tissues were re-fixed with formalin then embedded in paraffin wax and sectioned with 5μm thickness. Slides were deparaffinized, hydrated, treated with 3% H2O2 to block endogenous peroxidase then incubated in a steamer for 30 minutes in Target Retrieval Solution (Dako, #S1699, #S2367) for antigen retrieval. Sections were treated with Protein Block (Dako, #X090930-2) then incubated with primary F4/80 (Cell Signaling Technology, #70076) or Iba-1 (Abcam, #ab178846) antibodies at 4℃ overnight. On the next day slides were washed then incubated with secondary antibody (Cell Signaling Technology, #8114S) for 30 minutes. DAB Substrate Kit (Cell Marque, #957D-20) and Hematoxylin (Fisher HealthCare, #220-101) were used for staining and counterstaining. Images were scanned with P250 High Capacity Slide Scanner (3DHISTECH) and viewed with CaseViewer (v2.4, 3DHISTECH). For staining quantification, tumor areas were mapped using QuPath (v0.3.2) as regions of interest(ROIs), then the ROIs underwent cell detection and positive staining detection under same parameters across all slides to get the total number of cells, total number and percentage of positively stained cells. GraphPad Prism 9.0 was used to analyze the results. To test the hypothesis of more macrophage infiltration into TKO tumors, a one-sided T-test was used to compare positive cell percentage in TKO vs DKO samples.

Total RNA was isolated using the RNAeasy Mini Kit (Qiagen) and reverse transcription of total RNA was performed with SuperScript™ IV One-Step RT-PCR System (Thermo Fisher Scientific). Quantitative PCR was performed using the SYBR Green PCR Master Mix (Thermo Fisher Scientific), as previously described^14^. GAPDH was used as the internal control. GraphPad Prism 9.0 was used to analyze the results. PCR primers for *Adgre1* (F4/80) were: (F: ATTGCGGGATTCCTACACTATC; R: TTCACCACCTTCAGGTTTCTC). PCR primers for *Aif1* (Iba-1) were: (F: CCCACCTAGAGCTGAAGAGATTA, R: GATCTCTTGCCCAGCATCATT). To test the hypothesis of more macrophage infiltration into TKO tumors, a one-sided T-test was used to compare expression of macrophage markers in TKO vs DKO samples.

### Transcriptomic and survival analysis in clinical cohorts of OS and other cancers

We downloaded clinical and transcriptomic data from the NCI TARGET OS cohort using the TARGET data portal^47, 48^. We downloaded clinical and transcriptomic data from the Kuijjer 2012 cohort using the R2 genomics portal (accession “Kuijjer - 127 - vst - ilmnhwg6v2”)^49^. We filtered to keep only patients with both RNA-seq and survival data and no abnormally low TPM values, leading to 83 patients for the NCI TARGET OS and 84 for the Kuijjer et al cohort. To find homologs between mouse and human genes, we used the R package biomaRt^50^. To calculate expression scores for the TKO overexpressed genes, we used the module score method, which bins genes by their expression level, selects random genes from similar expression bins, calculates means of test and random genes, and subtracts the difference^51^. To calculate the “TKO myeloid only” signature, we used the MSIGDB C8 cell signature category downloaded via msigdbr and searched for all gene sets with the term “myeloid” in the title, then took the intersect of the genes in these sets with the significantly upregulated TKO genes. For deconvolution analysis of the NCI TARGET OS and Kuijjer 2012 cohort RNA-seq data, we used Microenvironment Cell Population Counter (MCP-Counter), the human counterpart of the mouse-focused mMCP-Counter.^52^

For survival analysis with the module scores and MCP counter infiltration scores, we used the survival (v3.4.0) and survminer (v0.4.9) packages in R for both Kaplan-Meier (KM) and Cox regression analysis. For KM we dichotomized the module scores or MCP-Counter scores at the median value to compare survival in high versus low with plots with KM and log-rank tests. For Cox regression, we input the scores as univariate continuous, dichotomized, and tercile analysis. Additionally, we performed the survival analysis on all clinical variables available in the NCI-TARGET data, including age, sex, and presence of metastases at diagnosis. We performed multivariable Cox regression for each module score and MCP-Counter using the only such clinical variable that was significantly associated with survival (metastasis at diagnosis). Finally, we assessed the assumption of proportional hazards for each Cox regression via graphical and statistical tests of Schoenfield residuals using the “ggcoxzph” and “cox.zph” functions and observed no deviations.

To test SKP2 correlations with other genes and with MCP-Counter immune infiltration scores, we used Pearson correlation with the cor.test function in R. For correlation analysis of all other genes, we also used multiple test correction with the false discovery rate (FDR) method and defined significant correlation as those with FDR < 0.05. We used Fisher exact tests to test for overlap of the TKO gene signature with the negatively correlated genes.

For TCGA analysis we downloaded data directly from the Genomic Data Commons website using the TCGABiolinks R package^53^. We used the cor.test function to test the correlation of SKP2 with MCP counter monocyte lineage score. We also used the Timer2.0 web portal to study the correlation of SKP2 expression and survival with monocyte and macrophage infiltration scores^54^.

## Results

### SKP2-KO induces overexpression of immune related genes

We recently developed a transgenic OS mouse model with germline *Skp2* knockout (“TKO”: Osx1-Cre;Rb1^lox/lox^;p53^lox/lox^;SKP2) and compared it to a baseline model (“DKO”: Osx1-Cre;Rb1^lox/lox^;p53^lox/lox^)^13^ for their pre-clinical pathologic features ^26^. The result showed that TKO mice experience improved survival compared to DKO mice, with delayed detectable tumorigenesis, slower tumor growth, increased apoptosis, and reduced stemness in TKO tumors.

To study the transcriptional effects of SKP2 in OS, we performed RNA-seq on OS tumors (n=3) from both models [**Fig 1A**]. We detected thousands of genes significantly differentially expressed with > 2 fold changes and at the adjusted p-value < 0.05 (1736 genes significantly upregulated, 940 downregulated) [**Fig 1B**]. SKP2 expression was confirmed to be completely abolished in the TKO samples [**Supplementary table 1**]. Surprisingly, many of the top genes overexpressed in the TKO were related to microenvironment signatures, especially immune-related genes including complement factors such as *C1qa* and major histocompatibility complex class 2 (MHC2) component *H2-Aa*. Gene set overrepresentation analysis showed that TKO tumors upregulated gene signatures related to immune function, muscle- and actin-related genes, endothelial function, and KRAS signaling [**Fig 1C, E**]. Concordant with these results, we observed significant upregulation in TKO of pan-immune marker *Ptprc* (CD45) and endothelial marker *Pecam1*, and modest upregulation of *Kras* [**Supplementary table 1**]. To infer what transcriptional networks and their top regulators are impacted in TKO tumors, we next analyzed enrichment of transcription factor targets and cancer-related pathway genes. Analysis of transcription factor target signatures from MSIGDB revealed upregulation of *MEF2A* targets, *CBFA2T2* and *CBFA2T3* targets, *TCF3* targets, and *TFAP4* targets, among others [**Fig 1E, middle panel, Supplementary Figure 1A**]. Additionally, we checked the enrichment of MSIGDB oncogenic signatures in the TKO upregulated genes [**Fig 1E lower panel, Supplementary Fig 1C].** We found significant enrichment of “SNF5_DN.V1_UP”, genes upregulated in *SNF5* knockout (also known as *Smarcb1* or *Baf47*), which was consistent with downregulation of *SNF5* targets in TKO. Furthermore, we observed upregulation of *MEK1* (*Map2k1*) and *Kras* targets.

**Figure 1:**
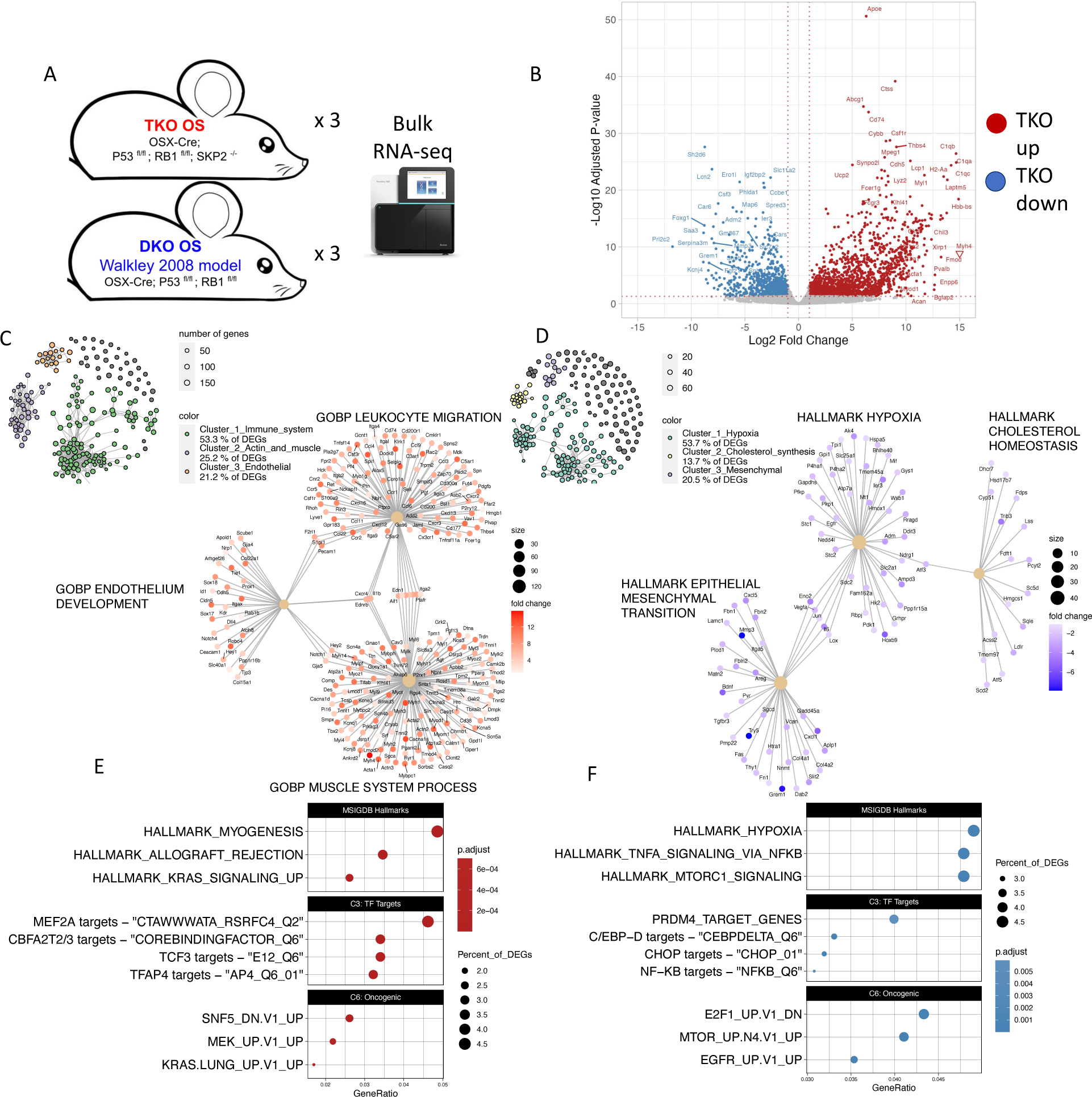
Differential RNA expression analysis. **A**): Experimental workflow. **B**): Volcano plot summarizing differential expression between TKO and DKO samples. **C,D**): Network-based gene set overrepresentation analysis of genes upregulated (C) or downregulated (D) in TKO. E,F): Dotplots for enrichment of MSIGDB Hallmarks and transcription factor targets, and oncogenic signals upregulated (E) or downregulated (F) in TKO.

Conversely, gene sets enriched among downregulated genes in TKO relative to DKO were dominated by signatures of hypoxia, along with lipid metabolism and EMT/invasiveness [**Fig 1D, F**]. Accordingly, we observed significant downregulation of hypoxia response factors *Hif1a* and vascular endothelial growth factor ligands *Vegfa* and *Vegfd*, cholesterol synthesis enzymes including *Sqle* and *Fdft1*, and extracellular matrix remodeling proteins such as *Mmp3* and *Try5* [**Supplementary table 1**]. Transcription factor activity downregulated by TKO included *PRDM4* targets, many of the C/EBP family targets including *C/EBPD* and *CHOP* targets, and *NF-KB* targets [**Fig 1F, middle panel, Supplementary figure 1B**]. Analysis of oncogenic signatures downregulated in TKO detected downregulation of the signature “E2F1_UP.V1_DN”, corresponding to genes downregulated after overexpression of *E2f1*. This is consistent with upregulation of *E2f1* targets in TKO. Additionally, we observed downregulation of *mTOR* and *EGFR* signatures [Fig 1F, lower panel, Supplementary figure 1D].

Taken together, the RNA-seq results suggest a critical shift of the tumor immune microenvironment in the TKO OS and link a set of critical downstream TFs and signaling that may mediate SKP2’s important roles in OS tumor development and progression.

### Deconvolution analysis reveals immune and stromal infiltration is enhanced in the TKO tumor microenvironment

The most parsimonious explanation for the increase in immune and other microenvironment signals in the TKO would be an increase in the infiltration of those cell types into the TKO microenvironment. Prior work on immune microenvironment gene expression in OS showed that immune-related gene expression was indeed driven by infiltrating immune cells, rather than by ectopic expression of immune genes in the malignant OS cells^36^.

Thus, we hypothesized that the level of immune cell infiltration would be greater in the TKO than in the DKO. To study this, we used a method of bulk RNA-seq deconvolution analysis called murine Microenvironment Cell Population Counter (mMCP-Counter)^44^. We chose this method because it is the newest such tool designed specifically for mouse tumor microenvironment deconvolution, rather than other well-known analogous tools like CIBERSORT, which are designed for humans. Applying it to our bulk RNA-seq samples, we observed that overall microenvironment infiltration levels were higher in the TKO compared to the DKO [**Fig 2A**]. However, this difference was driven by specific cell types, with the strongest enrichment observed for macrophage cells [**Fig 2B**]. Other significant differences included B cells, blood vessel-related cells including endothelial cells, T cells broadly and specifically CD8 T cells, and basophils. To corroborate this finding, we examined the expression of canonical markers of these cell types and observed significant differences in most of them [**Fig 2C**]. Markers of malignant OS cells or osteoclasts, which are bone-resident macrophage-like cells involved in bone resorption, were not significantly differentially expressed.

**Figure 2:**
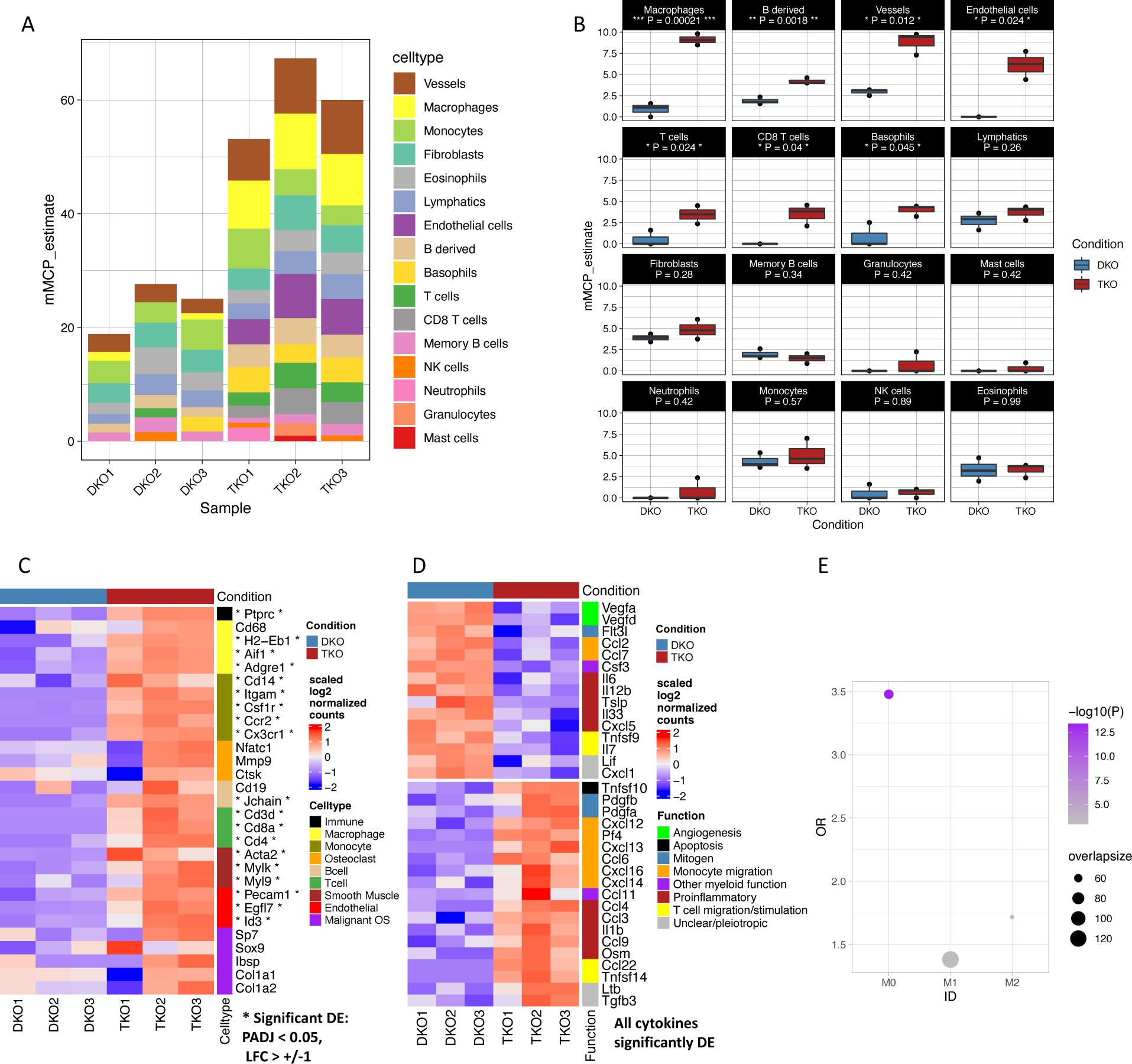
Microenvironment cell deconvolution analysis of bulk RNA-seq samples. **A**): Barplots summarizing mMCP-Counter results for all cell types in each sample. **B**): Boxplots illustrating differential estimates of enrichment between TKO vs DKO. Asterisks indicate statistical significance (t-test). **C**): Heatmap showing expression of specific markers of various infiltrating cell types. Asterisks indicate statistically significantly differentially expressed genes between TKO vs DKO. **D**): Heatmap showing expression of differentially expressed cytokines. **E**): Enrichment analysis of TKO upregulated genes among markers of M0, M1, or M2 macrophages.

Because immune infiltration and activity is often mediated by cytokine expression, we investigated the expression of all cytokine genes and found that many were significantly differentially expressed [**Fig 2D**], but unfortunately the data did not provide a clear pattern on which cytokines might play an active role in recruiting immune cells to TKO OS. For example, CCL2 is important for macrophage infiltration in many cancers but its expression was reduced in the TKO tumors^55, 56^. To address the potential functional impacts of increased macrophages in TKO OS, we also characterized the macrophage polarization status. We reanalyzed a previously published single-cell RNA-sequencing data from unstimulated “M0” macrophages, macrophages stimulated *in vitro* to an “M1” phenotype via exposure to lipopolysaccharides, and macrophages stimulated to “M2” phenotype via exposure to IL4 to define specific marker genes for these polarization states [**Supplementary figure 2**]^57^. We found that genes overexpressed in TKO showed significant enrichments for genes expressed higher in all three states, but the strongest enrichment was for genes related to the M0 phenotype [Fig 2E], suggesting that the macrophages in the TKO OS may be in a “naïve” state.

### Immune markers detected by RNA-seq are expressed higher in SKP2 KO tumors at the RNA and protein levels

We next used two independent assays to validate our RNA-seq findings of increased immune cell infiltration in TKO tumors. Because macrophages exhibited the greatest difference between TKO and DKO tumors, we focused on confirming their differential abundance. First, we applied quantitative RT-PCR (qPCR) to compare the expression of macrophage marker genes *Adgre1* (F4/80) and *Aif1* (Iba-1). We found that these genes were expressed at a higher level in the TKO tumors relative to DKO (P = 0.042 and P = 0.028, respectively) [**Fig 3A, B**]. We also observed a high variability in the expression of these markers among the TKO tumors.

**Figure 3:**
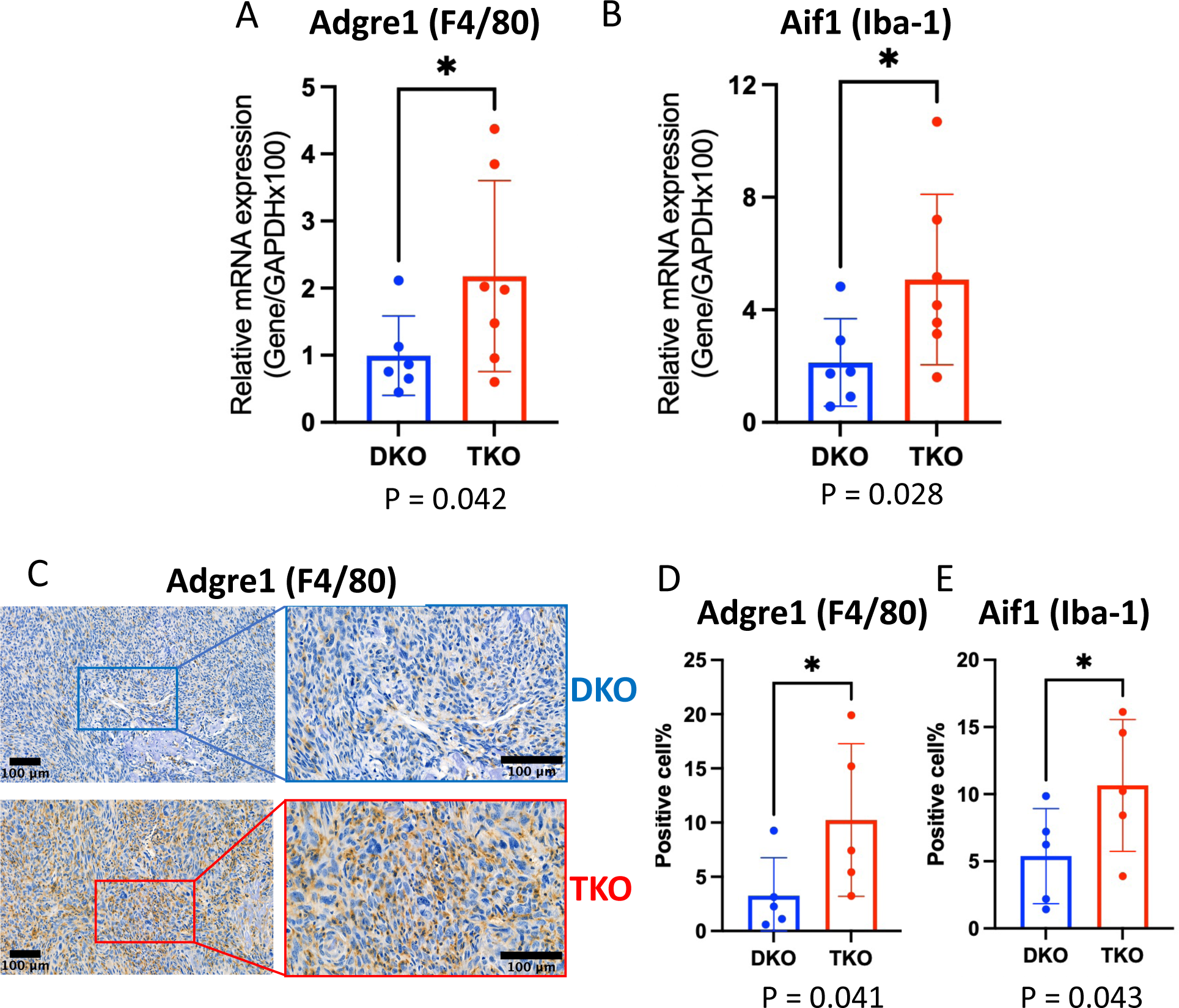
Validation of upregulation of macrophage markers in TKO. **A,B**): Quantitative PCR results for macrophage markers Adgre (F4/80) and Aif1 (Iba-1), respectively, in TKO vs DKO. **C**): Representative image of immunohistochemistry (IHC) for macrophage marker Adgre1 (F4/80) staining in TKO vs DKO tumors. **D,E**): Quantification IHC-derived positive cell percentage and comparison for Adgre (F4/80) and Aif1 (Iba-1), respectively, in TKO vs DKO.

Next, we performed quantitative analysis of immunohistochemistry staining for the same macrophage markers, *Adgre1* (F4/80) and *Aif1* (Iba-1). In general, we observed more positive cells in the TKO tumors compared to those from DKO [**Fig 3C**]. After semi-quantification of positive cell percentages, we found a higher percentage of cells positive for both F4/80 and Iba-1 in TKO relative to DKO tumors (P = 0.041 and P = 0.043, respectively) [**Fig 3D, E**]. As in the qPCR assays, we observed a high variability in the percentages of F4/80+ and Iba-1+ cells among the TKO tumors.

### Genes upregulated in SKP2 KO tumors are associated with improved survival in OS patients

We have previously found that TKO mice have significantly improved survival compared to DKO mice^26^. Additionally, we have shown that low expression of SKP2 was correlated with improved overall survival and metastasis-free survival in two distinct cohorts of OS patients^25^. To test if SKP2-related gene expression is associated with survival in OS patients, we performed survival analysis of the National Cancer Institute (NCI)’s TARGET OS cohort^48, 58^.

We hypothesized that patients highly expressing genes that were upregulated in the TKO samples would exhibit improved survival over patients with lower expression of those genes. After converting the mouse genes to their human homologs, we calculated a module score for all the genes that were either significantly up-regulated or down-regulated in the TKO samples, to obtain a score for the up- and a score for the down-genes for each patient. We observed a significantly improved 5-year overall survival in OS patients with high expression of the TKO upregulated genes, compared to the patients with low expression of the same genes [**Fig 4A**]. Since many of the up-regulated genes are immune related, we next investigated if the signal was driven by immune signatures. Using the Molecular Signatures Database (MSIGDB)^59^, we selected gene sets related to myeloid cell types from the “C8” cell type marker category, and then took the genes in those sets that were upregulated in the TKO. When we tested this TKO-upregulated myeloid immune signature, we found an even stronger survival benefit [**Fig 4B**]. Independently, we performed deconvolution analysis in the NCI TARGET data using Microenvironment Cell Population Counter (MCP-counter), the human counterpart of the mouse-specific mMCP-Counter method, then tested the survival association of infiltrating immune cell abundance^60^. High MCP infiltration scores for macrophages and CD8+ T cells were significantly associated with improved survival [**Fig 4C, 4D**]. Additionally, to account for potential confounding factors, we applied independent multivariable Cox models on the MCP-Counter scores using metastasis at diagnosis as a covariate and confirmed the survival benefits for each score [**Supplementary table 2**].

**Figure 4:**
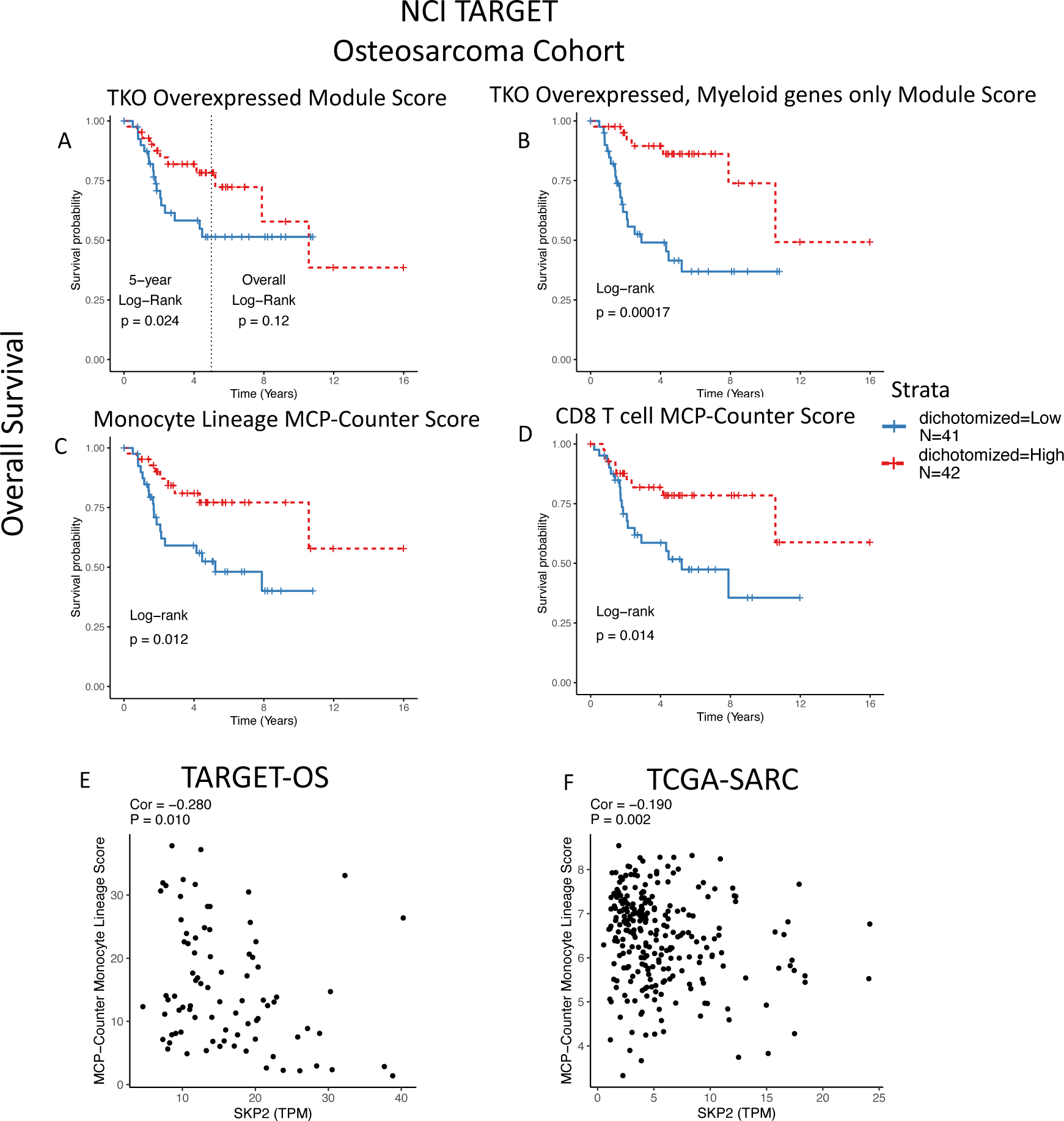
Correlations of up-regulated genes in SKP2 KO tumors with macrophages and improved survival in patients. **A-D**): Kaplan-Meier (KM) plots showing overall survival for patients from the NCI TARGET OS cohort with high versus low scores for the TKO upregulated gene module; the module of myeloid-only genes from the TKO upregulated signature; MCP-Counter monocyte lineage infiltration score; and MCP-Counter CD8 T cell score, respectively. For each variable, the cohort was dichotomized at the median to compare survival in high-score vs low-score patients. **E, F**): SKP2 correlation analysis with MCP-Counter monocyte lineage score for the TARGET OS and TCGA-Sarcoma cohorts, respectively.

We attempted to validate our findings in a separate cohort of OS patients. We used data from a 2012 study in which 84 participants donated pre-treatment biopsy samples for microarray gene expression analysis, downloaded from the R2 genomics database (the “R2 Kuijjer cohort”)^49^. In this cohort, we found modest but consistent associations with improved metastasis-free survival for the TKO signature and MCP macrophage and T cell infiltration scores [**Supplementary figure 4 A-D**].

Taken together, these results indicate a positive association with patient survival for the genes upregulated in the SKP2 KO and the association is likely explained by increased macrophage infiltration to the OS.

### SKP2 is negatively correlated with macrophage infiltration in OS and other sarcoma patients

To find genes and pathways associated with SKP2 expression in the human OS tumors and how they may be related to macrophage infiltration, we next performed two types of correlation analysis in the NCI-TARGET OS cohort. First and remarkably, we found that SKP2 expression itself in the TARGET-OS patients was significantly negatively correlated with macrophage infiltration scores as estimated by MCP-counter [**Fig 4E**]. In the Kuijjer cohort, we also observed a significant negative correlation between SKP2 expression and monocyte infiltration scores (r = −0.48, P < 4e-6) [**Supplementary Figure 4E**]. This is consistent with our finding of increased macrophages in our Skp2 KO mouse model (i.e., reduced macrophages in the Skp2 WT OS).

To identify genes that may be regulated by SKP2 in the patient OS tumors, we searched for genes whose expression is significantly correlated with SKP2 expression in the patients. We identified hundreds of genes significantly correlated with SKP2 expression in the NCI target OS cohort [**Supplementary table 3**]. The positively correlated genes were enriched in pathways related to cancer progression, mRNA processing, and cell proliferation. Remarkably, in the negatively correlated genes, we observed pathway enrichment primarily related to immune function, especially macrophages [**Supplementary figure 4 B,D**]. Furthermore, we found that the TKO overexpressed genes were significantly enriched in the negatively correlated genes (odds ratio = 3.95, P < 0.001; Fisher’s exact test).

Finally, to test if the correlation between SKP2 expression and macrophages also exists in other cancer types, we extended our analysis to all the tumor samples in The Cancer Genome Atlas (TCGA). This pan-cancer analysis showed that SKP2 was significantly and negatively correlated with MCP-Counter macrophage infiltration scores in the TCGA-SARC cohort, which is composed of 260 soft-tissue sarcoma patients [**Fig 4G**], but not other cancers. To strengthen our finding, we expanded our study using the Timer 2.0 web portal, which applied multiple software to estimate immune cell abundance in the TCGA tumors [**Supplementary figure 5**]^54^. The TCGA Sarcoma cohort consistently showed negative correlations between SKP2 expression and monocyte/macrophage infiltration scores estimated by multiple methods, as well as consistent survival benefits of macrophage infiltration. Other cancer types also showed some evidence of negative correlation of SKP2 with monocyte/macrophage infiltration or survival association with infiltration, but the results were less consistent than what was obtained for the TCGA-Sarcoma.

## Discussion

We have previously shown that SKP2 knockout drives improved survival and prognosis in pre-clinical models of OS^26^. Here, we show that *SKP2* knockout leads to a dramatic remodeling of the tumor microenvironment. Multiple cell types including macrophages, T cells, B cells, and vascular cells preferentially infiltrate into the *SKP2*-KO tumors, compared to the *SKP2*-intact tumors. To our knowledge, no prior connection of *SKP2* with any solid tumor malignancy tumor microenvironment has been reported, and this is the first high throughput transcriptomic assay performed in the context of *SKP2* knockout.

We demonstrated that high expression of the genes upregulated in the *SKP2*-KO OS tumors was correlated with improved survival in OS patients. This seems to be explained by greater expression of macrophage-related genes in the *SKP2* KO tumors. This is consistent with a previous report of the positive impact of macrophages on prognosis in OS^36^. Strikingly, we discovered a consistent negative correlation between SKP2 expression and macrophage infiltration in patient cohorts of OS and other cancer types including soft-tissue sarcomas, suggesting that regulation of the tumor microenvironment could be a key function of SKP2 that extends beyond pre-clinical mouse models. This is especially relevant in the context of recent reports indicating that antibody-based blockade of immunomodulatory signals Cd47 and Gd2 drove macrophage-mediated anti-tumor cytotoxicity via phagocytosis in OS^37^. Additionally, another recent study also indicated that inhibition of L-amino acid transporter 2 (LAT2) enhanced anti-tumor macrophage immunity and sensitized OS to chemotherapy^61^. Taken together, these studies provide a rationale for testing potent future combination therapy strategies in OS that aim to promote immune recruitment via *SKP2* inhibition and promote macrophage activation via anti-CD47/GD2 or anti-LAT2 therapeutics.

The precise mechanism underlying the remodeling of the TME in relation to SKP2 is beyond the scope of the current study. Nevertheless, we detected differential expression of thousands of genes, among which are cytokines, transcription factors, and known cancer-associated genes which may explain our observations. Many of these factors are critical to the regulation of cell proliferation, cell differentiation, and immunity. Therefore, their disruptions may help explain both the improved survival and enhanced immune infiltration in SKP2 KO tumors, but also help uncover the resistance mechanism employed by OS to *SKP2* modulation. Several of the transcription factors upregulated upon SKP2 KO have previously been reported as proteolytic targets SCF^SKP2^ ubiquitination, such as *TCF3* or Mef2 family members *MEF2C* and *MEF2D*^62, 63^. We also observed evidence for *E2F1* upregulation upon SKP2 KO, which has a major effector of apoptosis upon SKP2 knockout in the context of RB1/P52 deficiency^18^. Furthermore, we observed downregulation of PI3K/mTOR/AKT pathway targets upon *SKP2* knockout, consistent with SKP2’s reported role of stabilizing AKT1 via protective K63 ubiquitination^64^.

In addition to such *SKP2* targets reported in literature, we also observed significant differential expression of the targets of other transcription factors for which no link with SKP2 has been reported to our knowledge, including upregulation of targets of the paralogs *CBFA2T2*, *CBFA2T3* along with *TFAP4*, and downregulation of *PRDM4* targets. Interestingly, TFAP4 was reported to undergo proteolytic ubiquitination via a different F-box protein related to SKP2 called BTRCP in conjunction with the SCF E3 ligase complex^65^. We also observed downregulation of targets of various CCAAT-enhancer-binding proteins (C/EBP) family members including *C/EBP-B*, *C/EBP-D*, and *CHOP*, as well as *NF-KB*. *NF-KB* and one C/EBP paralog, *C/EBP-A*, were reported to be inhibited by SKP2 and thus many of their targets would be expected to increase expression in the SKP2 KO tumors; however, we found that many of them actually decreased expression in TKO^66, 67^. Paradoxically, many genes involved in growth factor signal transduction were differentially expressed in a discordant manner, including upregulation of KRAS and MEK1 (Map2k1) targets and downregulation of EGFR in SKP2 KO tumors. It is possible that previously described regulatory relationships may be altered *in vivo* in the context of Rb1/P53 doubly deficient OS. Ultimately, the interaction among different signals and pathways will need more studies in the future in order to fully understand the benefits and caveats of targeting SKP2.

There are limitations in current study. One is that the technical nature of bulk RNA-seq does not allow direct profiling of the gene expression in individual cellular components. This may have limited our ability to address if the OS malignant cells directly promote the recruitment of macrophages to tumors, for examples, our comparison of cytokine analysis was based on the cytokines produced by all cell types. New methods like single-cell RNA-sequencing or spatial transcriptomics will allow more precise study of the effects of SKP2 KO in distinct populations of malignant cells and TME compartments. Additionally, it is unclear if SKP2 exerts the described microenvironment-remodeling effect within malignant cells, infiltrating immune cells, or both, due to the organism-wide nature of the germline SKP2 knockout. While this model does reflect the clinical setting of system-wide SKP2 inhibition, a secondary model of conditional SKP2-KO via OSX-Cre lox would allow us to study the SKP2-monocyte recruitment mechanism in more details. Lastly, considering the genetic diversity of OS and the variability of our models, additional samples would help strengthen our results.

## Supporting information

Supplemental material folder containing supplement figures and tables

## Author Contributions

A.F. performed bioinformatic and clinical data analysis. J.W. performed tumor isolation and RNA-seq analysis. R.Z. performed RT-PCR and IHC assays. B.K.F assisted with IHC quantification and analysis. W.A.H, S.S., and H.B. assisted with mouse model preparation and tumor isolation. E.S. and H.Z. assisted with data interpretation. R.Y. and D.G. assisted with clinical data analysis and data interpretation. B.H. and D.Z. conceived the study, supervised all work, and acquired funding. D.Z. supervised all data analysis. A.F., B.H. and D.Z. wrote the manuscript with inputs from all authors.

## Data availability

Data has been deposited to Gene Expression Omnibus (GEO).

