## Supplemental material folder containing supplement figures and tables for "SKP2 knockout in Rb1/p53 deficient mouse models of osteosarcoma induces immune infiltration and drives a transcriptional program with a favorable prognosis": Ferrena_Supplemental_Figures.pdf

**Supplementary Tables 1-3 in a separated Excel file**

**Supplementary Table 1:** DESeq2 results and DESEq2 size-factor normalized counts for TKO vs DKO bulk RNAseq samples.

**Supplementary Table 2.** Multivariable model results from four Cox models, where each variable was loaded with Metastasis at diagnosis as a covariable.

**Supplementary Table 3:** SKP2 gene correlation results from NCI TARGET OS cohort.

**Supplementary Figure 1: transcription factor and oncogene signature overrepresentation analysis in TKO vs DKO. A,B**): Top panels: Overrepresentation analysis of transcription factor targets upregulated (A) or downregulated (B) in TKO, respectively. Lower panels: Gene-pathway dotplots for Fig 1E and 1F transcription factor targets. **C,D**): Overrepresentation analysis of oncogene signal targets upregulated (C) or downregulated (D) in TKO, respectively. Lower panels: Gene-pathway dotplots for Fig 1E and 1F oncogene signal targets.

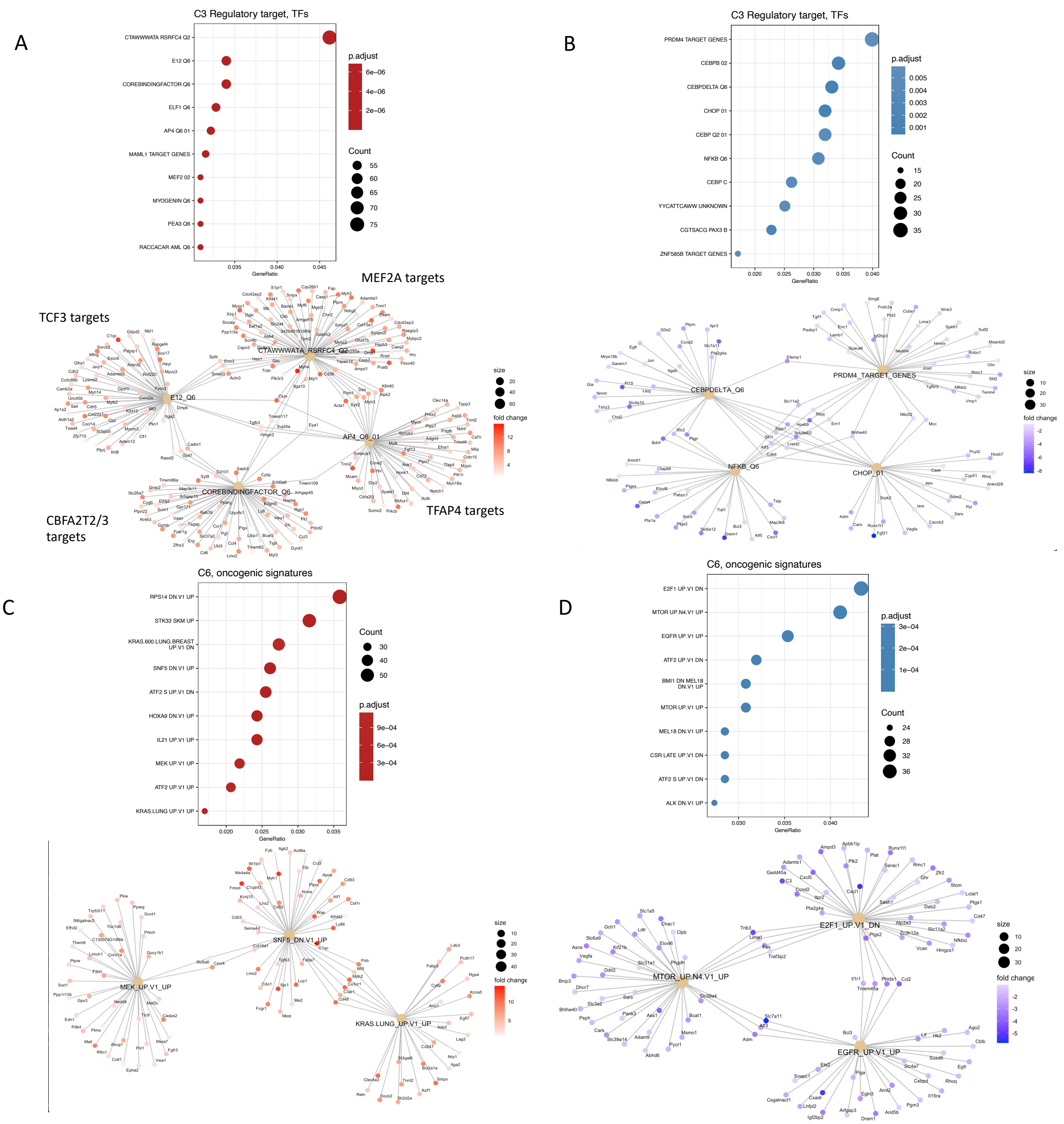

**Supplementary Figure 2: macrophage polarization marker derivation from reanalysis of MacSpectrum single-cell RNA-seq data. A**): integration result of MacSpectrum data with integration software RISC, visualized as a Uniform Manifold Approximation Projecton (UMAP) plot. **B**): Violin plots showing distribution of module scores calculated from detected markers of M0, M1 and M2 to check specificity of the markers. **C**): Heatmap of top 20 markers for M0, M1, and M2 derived from marker analysis of integrated data via Wilcoxon tests. **D**): All significant markers from M0, M1, and M2, plotted with gene-clustering to check specificity of markers.

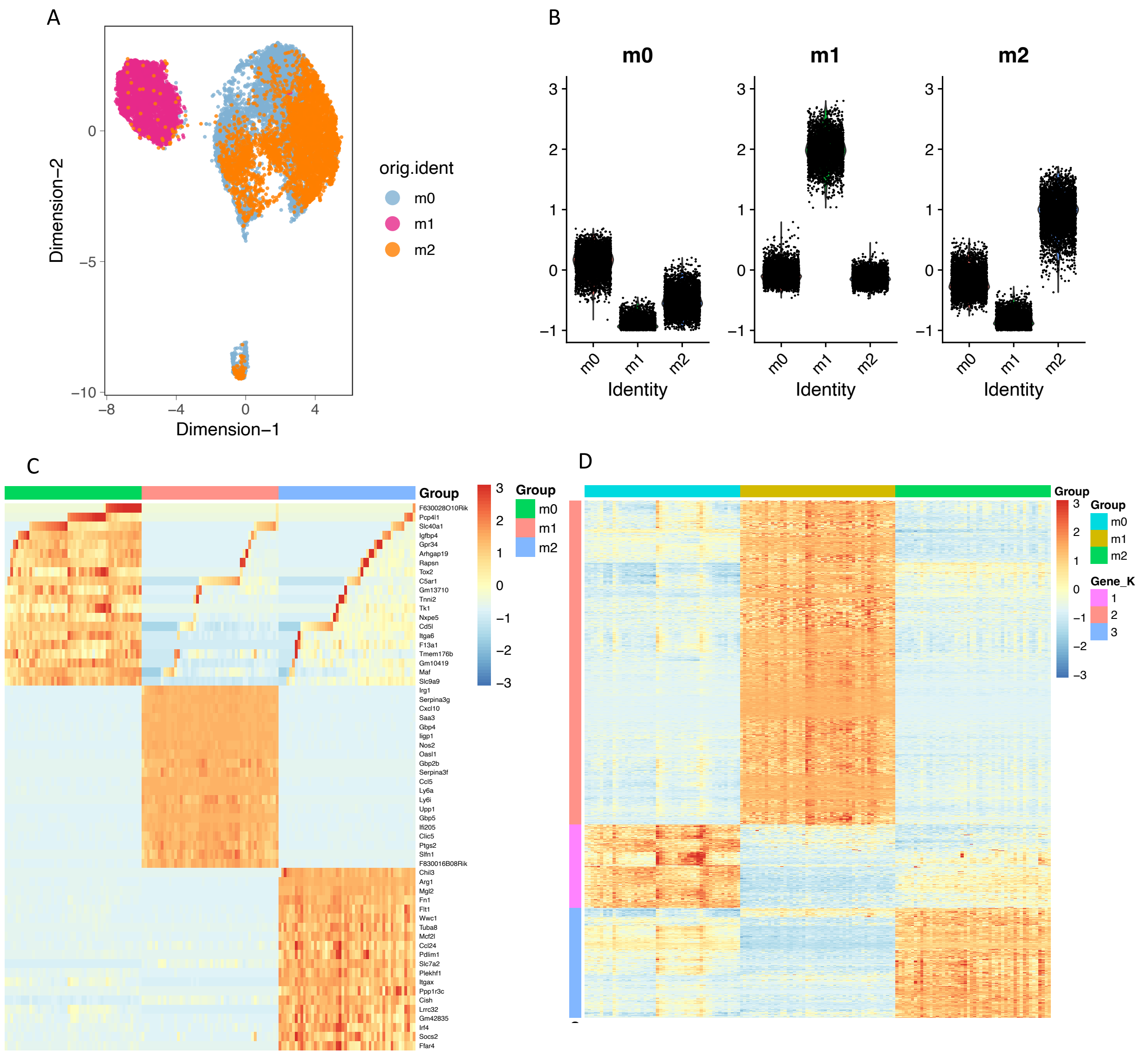

**Supplementary Figure 3: SKP2 KO signature is associated with macrophages and improved survival in Kuijjer 2012 OS cohort. A-D**): Kaplan-Meier (KM) plots showing metastasis-free survival for patients from the Kuijjer 2012 cohort with high versus low scores for the TKO upregulated gene module; the module of myeloid-only genes from the TKO upregulated signature; MCP-Counter monocyte lineage infiltration score; and MCP-Counter CD8 T cell score, respectively. For each variable, the cohort was dichotomized at the median to compare survival in high-score vs low-score patients. **E**): SKP2 correlation analysis with MCP-Counter monocyte lineage score for the Kuijjer 2012 cohort.

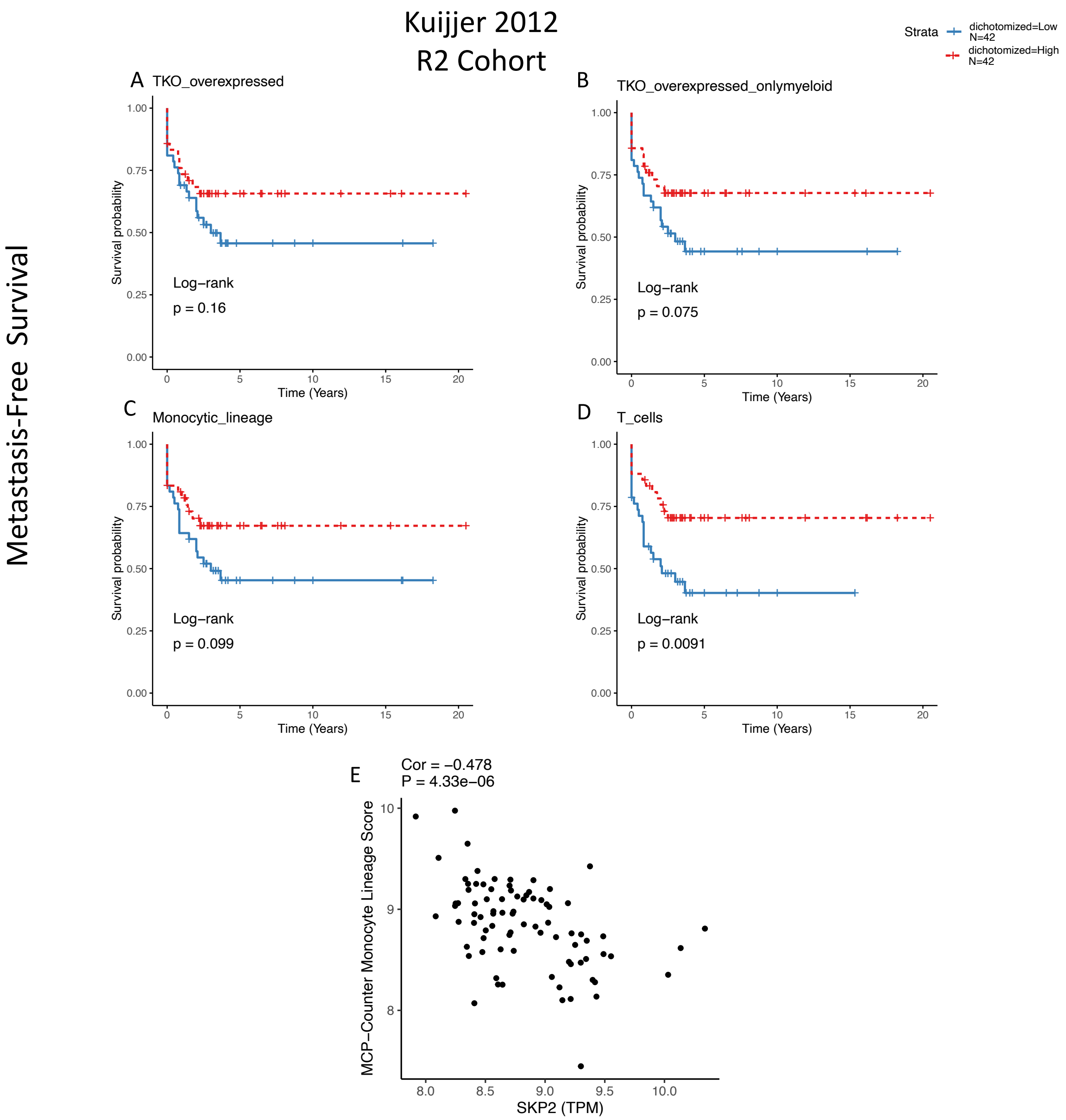

**Supplementary Figure 4: Gene set overrepresentation analysis of genes significantly correlated with SKP2 in the NCI TARGET OS cohort. A,B**): dotplots showing significant enrichment of gene sets derived from the entire MSIGDB database in positive and negatively correlated genes, respectively. **C,D**): dotplots showing significant enrichment of gene sets derived from the Gene Ontology Biological Process database in positive and negatively correlated genes, respectively.

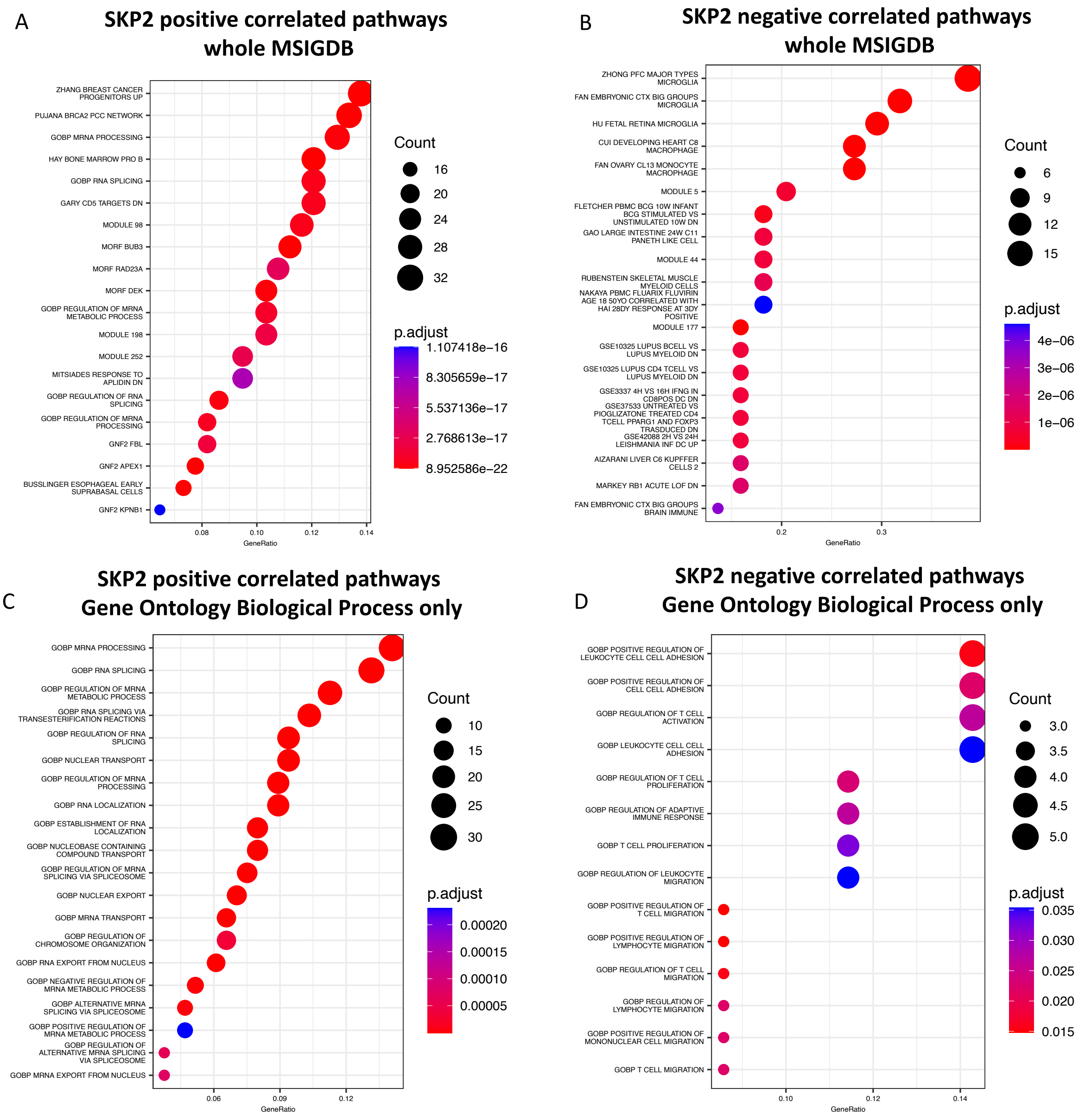

**Supplementary Figure 5: Pan-cancer TCGA analysis of macrophage infiltration correlation with SKP2 and survival via TIMER2.0. A,B**): SKP2 expression correlation with monocyte infiltration scores and macrophage infiltration scores, respectively, as measured by a variety of deconvolution methods, across TCGA cohorts. **C,D**): Survival analysis of monocyte infiltration scores and macrophage infiltration scores, respectively.

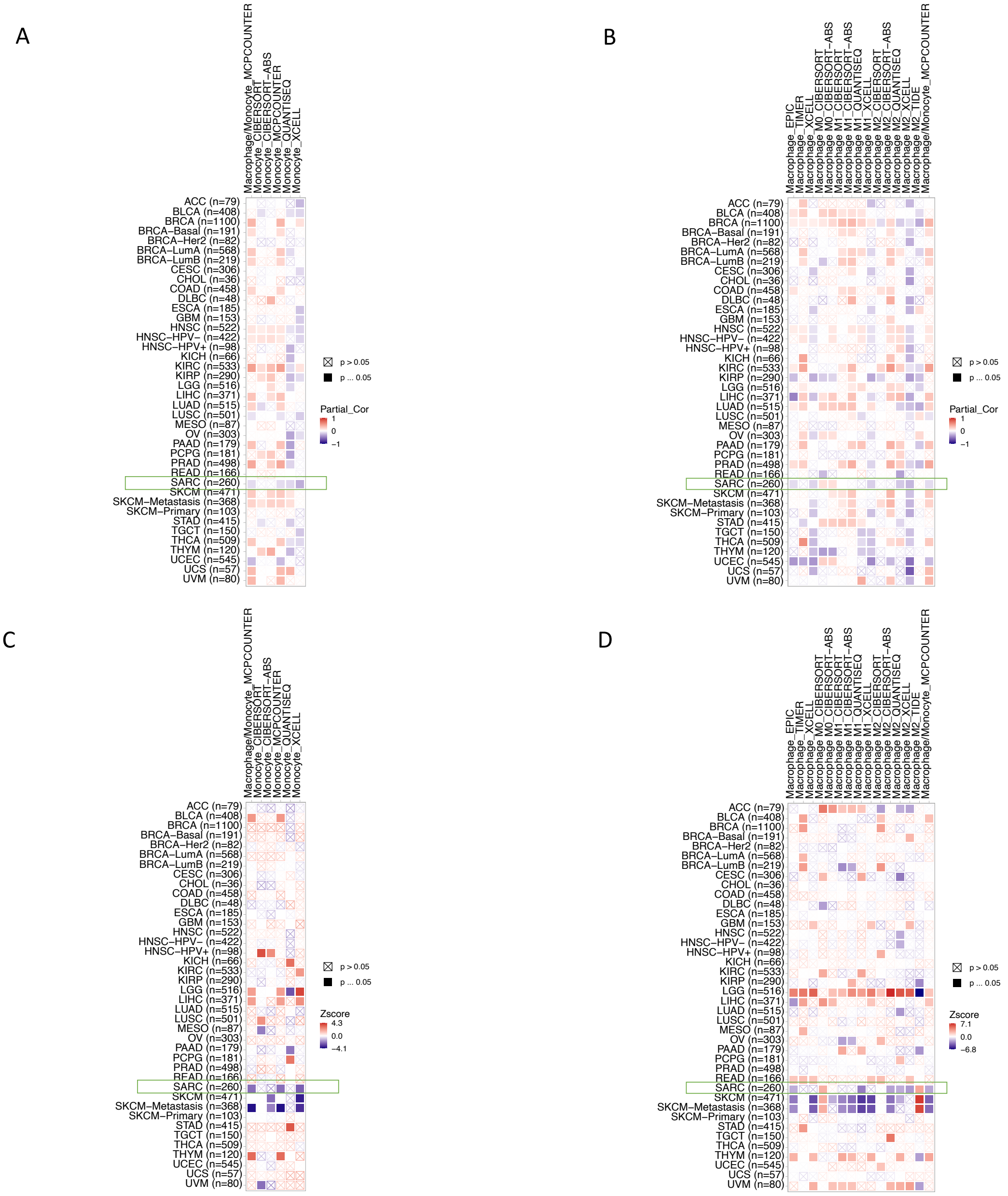
